# IBRAP: Integrated Benchmarking Single-cell RNA-sequencing Analytical Pipeline

**DOI:** 10.1101/2022.09.26.509481

**Authors:** Connor H. Knight, Faraz Khan, Upkar Gill, Jun Wang

**Author notes:** Corresponding author: Jun Wang, Centre for Cancer Genomics and Computational Biology, Barts Cancer Institute, Queen Mary University of London, London EC1M 6BQ, United Kingdom.

## Abstract

Single-cell RNA-sequencing (scRNA-seq) is a powerful tool to study cellular heterogeneity. The high dimensional data generated from this technology are complex and require specialised expertise for analysis and interpretation. The core of scRNA-seq data analysis contains several key analytical steps, which include pre-processing, QC, normalisation, dimensionality reduction, integration, and clustering. Each step often has many algorithms developed with varied underlying assumptions and implications. With such a diverse choice of tools available, benchmarking analyses have compared their performances and demonstrated that tools differentially operate according to the data types and complexity. Here, we present Integrated Benchmarking scRNA-seq Analytical Pipeline (IBRAP) – a tool, which contains a range of analytical components that can be interchanged throughout the pipeline alongside multiple benchmarking metrics that enables users to compare results and determine the optimal pipeline combinations for their data. We apply IBRAP to single and multi-sample integration analysis using pancreas, cell line and simulated data accompanied with ground truth cell labels, demonstrating the interchangeable and benchmarking functionality of IBRAP. Our results confirm that the optimal pipelines are dependant of individual samples and studies, further supporting the rationale and necessity of our tool. We then compare reference-based cell annotation with unsupervised analysis, both included in IBRAP, and demonstrate the superiority of the reference-based method in identifying robust major and minor cell types. Thus, IBRAP presents a valuable tool to integrate multiple samples and studies to create reference maps of normal and diseased tissues, facilitating novel biological discovery using the vast volume of scRNA-seq data available.

## Introduction

Single-cell RNA-sequencing (scRNA-seq) has innovated transcriptome characterisation, enabling high-resolution investigations into heterogenous cell populations in health and disease. A variety of library preparation procedures are commercially available, broadly categorised as plate- and droplet-based methods, distinguished by different components in the infrastructure, such as, transcript coverage, cell sorting, unique molecular identifier inclusion, and more [1]. Regardless, all platforms downstream result in files that align against transcriptomes with alignment software that generate count matrices representing individual cells, genes, and the number of corresponding transcripts. Post-alignment bioinformatic processing predominantly encompasses the following stages: pre-processing, QC, normalisation, integration, clustering, and differential expression analysis (DEA) [2]. Numerous analytical packages and components are available for each of these steps. This is evident by the fact that scRNA-seq packages have increased from 84 (01/01/2017) to 1,192 different packages (18/03/2022) that are available largely spread across Python (39.7%) and R (58.1%). With numerous algorithm choices, it is difficult for community members to decide which application is best for their pipeline, especially considering that each algorithm has its own set of strength and limitations [3].

Several benchmarking studies have been performed for either individual aspects of scRNA-seq bioinformatics (i.e. normalisation, integration, clustering) or as pipelines as a whole. Conclusively, there was no single approach able to gain the best results across all sample scenarios which leads to multiple tool recommendations or case-by-case recommendations [2,4–12]. Since tools are not integrated, custom pipeline coding is required which is tedious with inevitable room for errors. Thus, there is a need for a tool that already combines different pipelines and tools, and capable of distinguishing pipeline performance.

Other well-established pipelines, including Seurat [13], Scanpy [14], and Pegasus [15] lack a variety of methodology for users to interchange, leaving the users with relatively linear approaches to their analyses (see Table 1). ASAP [16], a web-based analytical pipeline sought to integrate a range of analytical methods, but is missing a large variety of single-cell-specific methods for single-cell data; a further limitation of ASAP is that that it lacks a variety of benchmarking capabilities to distinguish performances.

**Table 1.** | Table of leading scRNA-seq pipelines for comparison to IBRAP. ASW = Average Silhouette Width, Adjusted Rand Index = ARI, Normalised Mutual Information = NMI.

|  | Droplet<br>Pre-processing | Normalisation | Integration | Reduction | Clustering | Trajectory<br>Inference | Benchmarking | Platform |
| --- | --- | --- | --- | --- | --- | --- | --- | --- |
| IBRAP | Scrublet,<br>DecontX | Seurat<br>(CPM), TPM,<br>Scran,<br>SCTransform | BBKNN,<br>Scanorama,<br>Seurat<br>CCA,<br>Harmony | PCA,<br>Diffusion<br>Maps, t-SNE,<br>UMAP,<br>LargeVis | kmeans, PAM,<br>SC3, louvain,<br>louvain with<br>multiplelevel<br>refinement, smart<br>local moving,<br>leiden, SingleR | Slingshot | ASW<br>(biological &<br>Batch), ARI,<br>NMI,<br>connectivity,<br>Dunn Index | R |
| ASAP | None | CPM,<br>DESeq2,<br>voom log2,<br>SCTransform | ComBat,<br>fastMNN,<br>Limma | PCA,<br>UMAP, t-SNE | kmeans, SC3,<br>louvain, louvain<br>with multilevel<br>refinement, smart<br>local moving,<br>leiden | None | No | Web |
| scCancer | SoupX,<br>scds | lognorm | Seurat<br>CCA,<br>Harmony,<br>Liger | PCA, t-SNE,<br>UMAP | Louvain | None | No | R |
| Scanpy | Scrublet | CPM | ComBat,<br>BBKNN,<br>scanorama,<br>MNN | PCA,<br>Diffusion<br>Maps, t-SNE,<br>UMAP | louvain, leiden,<br>Ingest | PAGA | No | Python |
| Seurat | None | lognorm,<br>SCTransform | Seurat CCA | PCA, ICA,<br>t-SNE,<br>UMAP | louvain, louvain<br>with multilevel<br>refinement, smart<br>local moving,<br>leiden | None | No | R |
| Pegasus/Cumulus | None | Counts per<br>100,000 | Harmony,<br>Scanorama | PCA,<br>Diffusion<br>Maps, FLE | louvain, leiden | FLE-based | kBET, kSIM | Python/web |

Here we present IBRAP, a wrapper package that contains unique combinations of several pipelines by interchanging normalisation (the number of methods, n=4), integration (n=4), and clustering (n=7) components to create a unique workflow, which includes pre-processing, QC, normalisation, integration, clustering and DEA in a dataset specific manner to maximise sensitivity and specificity of identifying the accurate cell (sub-)types. Furthermore, we include 5 benchmarking metrics that can be applied to distinguish clustering performance and a metric for users to assess batch effect removal. We enlisted 15 publicly available and generated 4 simulated datasets using splatter [17] and applied all possible pipelines (adjusting parameter changes for expected clusters), to demonstrate the functionality of IBRAP and benchmark individual and integrated analysis. We further investigated the reference-based cell annotation compared with the unsupervised analysis in their ability to identify robust major and rare cell types, and demonstrated the potential usage of IBRAP in a reference map or Atlas like projects.

## Materials and Methods

### Workflow Overview

IBRAP consists of four main modules, namely data transformation, inference, Rshiny application and biological analysis (Figure 1). Within the data transformation module, data QC is performed, including doublet removal and ambient RNA decontamination (if needed) followed by normalisation, highly variable gene (HVG) selection and dimensionality reduction. Within the normalisation step, four leading methods are included, including Scanpy (CPM) [14], SCTransform [18], Scran [19], and TPM. For dimensionality reduction, five different techniques, PCA [14,20], Diffusion Maps [21], t-SNE [22], Largevis [22], and UMAP [18], were incorporated (Table 1). In addition to single sample analysis, IBRAP also incorporated four commonly used multi-sample integration methods that showed good performances in benchmarking studies, including BBKNN [23], Harmony [24], Seurat CCA [18], and Scanorma [25] which allows users to perform integration and compare performances for their samples.

**Figure 1.**
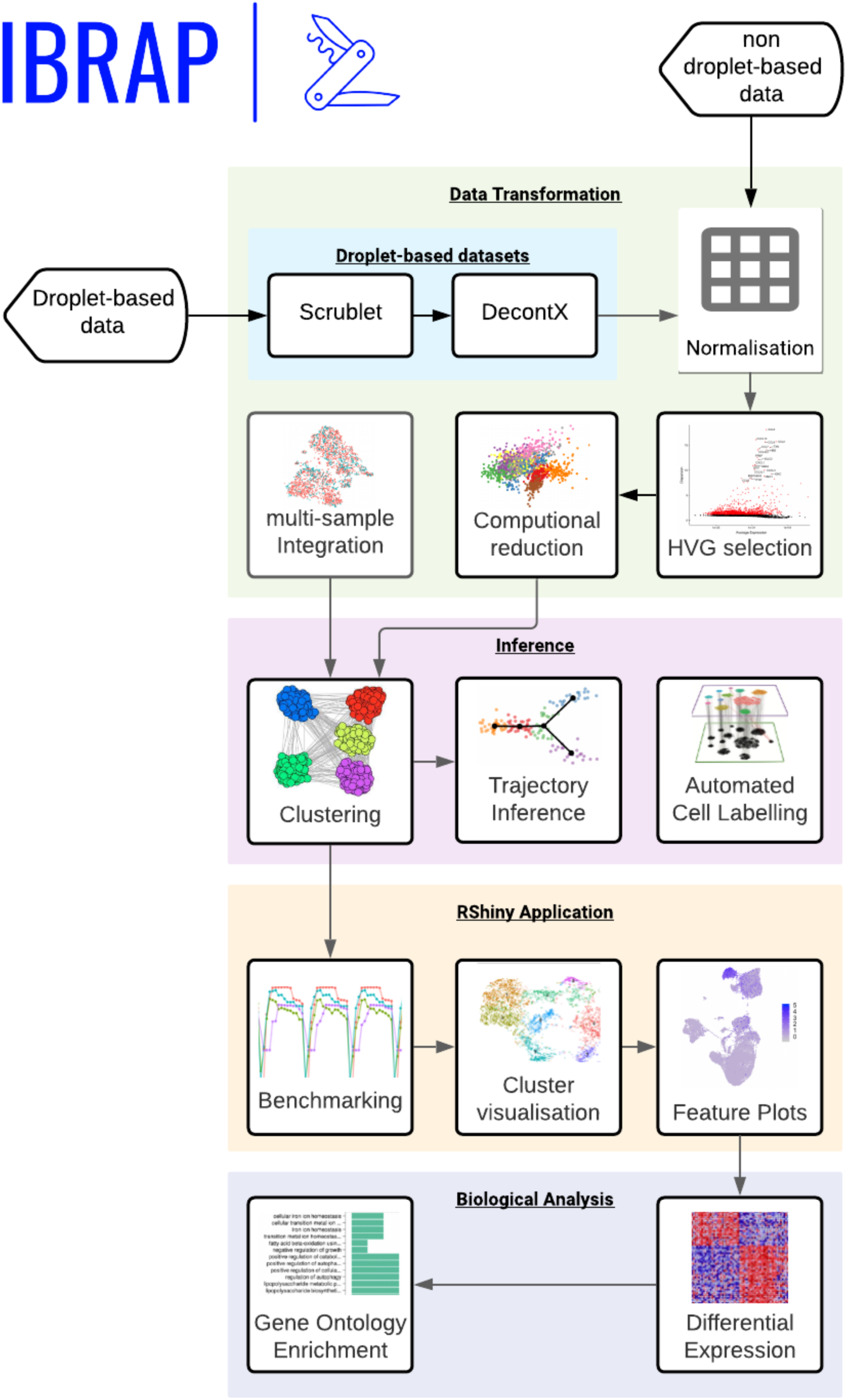
| A workflow for single-cell RNA-sequencing analyses in IBRAP. IBRAP accepts droplet- and non-droplet-based scRNA-seq counts. If the user has processed their cells with droplet-based infrastructure, they may use our droplet-based cleaning packages that we included (DecontX and Scrublet). Otherwise, the user may continue to data transformation encompassed with normalisation, highly variable gene selection, and when required, sample integration. After this has been performed a user then proceeds to inference, where a user will identify cell clusters, identify developmental trajectories, or label their cell types using a reference-based package. The user can then use our Rshiny application to investigate their results in an easier fashion than using the terminal. Finally, if the user opted for finding cell clusters, they must uncover the biology driving the clusters to identify their cell type. For this, the user can produce a range of different gene expression plots, differential expression, and a gene ontology analysis.

Within the inference module, IBRAP is able to perform clustering, trajectory inference and automated cell labelling (Figure 1). Within the clustering step, a selection of popular clustering techniques was integrated, including k-means, PAM, SC3, Louvain, Louvain with Multilevel Refinement, Smart Local Moving, and Leiden. SingleR [26], was also included in the inference module which allows users to perform supervised clustering based on a reference dataset to achieve automated cell annotation. By allowing the interchange of a diverse selection of methods within key analytic steps, e.g., normalisation, integration, reduction and clustering, IBRAP is able to create a large number of method combinations (pipelines) and subsequent results. Indeed, IBRAP contains up to 4 normalisation, 5 visualisation, and 7 clustering methods for individual samples and 4 integration methods for integration analyses. Given the large number of method combinations (i.e., pipelines) produced, we created the benchmarking function within IBRAP that produces and visualises benchmarking matrices and distinguish optimal results for individual samples and studies. Consequently, IBRAP produces a large volume of data, hence we designed object orientated functions around a novel S4 class object.

### Case Study Samples

#### Pancreas

All 8 datasets (cells=14,890) were available from the package SeuratData containing 13 cell type ground truth labels. Datasets were derived from 4 individual studies [27–30] using 5 technologies: CELseq, CELseq2, SmartSeq2, Fluidigmc1, and InDrop. Further information can be seen in Supplementary Table 2.

#### Cell Lines

Seven mixed cell line samples (cells=6,228) were acquired for a previously performed mixture control study [31]. Three of these samples contain cell lines: HCC827, H1975, and H2228, and four samples contain cell lines: HCC827, H1975, H2228, H838, and A549. These samples were processed with either: 10x Genomics, Dropseq, CELseq2. Further information can be seen in Supplementary Table 2.

#### Simulations

Four simulation datasets (cells=4,000) were created using the R package, Splatter. Each sample contained 20,000 genes. A total of 4 samples were produced: (1) 4 cell types and the number of cells was split equally amongst the clusters, (2) 4 cell types and the number of cells were unequally distributed amongst the clusters, (3) 8 cell types and the number of cells were equally spread amongst clusters, and (4) 8 cell types and the number of cells were unequally spread amongst clusters. Other parameter settings deviated from the default including: mean.shape=0.5, de.prob=0.2, lib.loc=12,lib.scale=0.7. Further information can be seen in Supplementary Table 2.

#### Core Analysis

We performed three analyses, one analysing the 19 datasets individually, a second integrating datasets from the pancreas and cell line cohort, and a third using supervised cell state annotation using references and cell type labels on query datasets. All analyses were used with IBRAP’s framework and closely followed default settings. In terms of filtration, cells were removed from the datasets if they contained over a calculated number of genes:

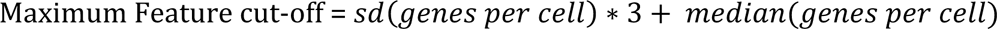

We also removed cells with less than 200 genes and when mitochondrial genes were detected, any cells containing over an 8% fraction. Any genes that were expressed in less than 3 cells were removed.

For the individual analyses, we implemented SCTransform, Scran, and Scanpy for normalisation methods with default parameters, regressing out total counts with Scanpy and Scran (note we did not do this for SCTransform since this is already included in the algorithm). We applied graph-based clustering algorithms: Louvain, Louvain with multilevel refinement (louvainMLR), smart local moving (SLM), and the Leiden algorithms; we also applied older clustering methods including kmeans, PAM, and SC3. For all clustering methods, we adjusted cluster diversity by adjusting either the resolution or expected clusters parameters. Following the single sample analysis, we next performed multi-sample integration using the same normalisation and clustering methods as the individual samples but using Seurat CCA, BBKNN, Harmony, and Scanorama to combine samples from different studies. Finally, to distinguish pipeline performance, we employed Average Silhouette Width (ASW), Adjusted Rand Index (ARI), and Normalised Mutual Information (NMI) benchmarking metrics. ASW compares the distance of cell labels in a UMAP projection to assess separateness, and ARI and NMI compare clustered cell labels against a ground truth list. We summarised these scores by ranking them individually, taking the average rank and then scaling from 0-100, 0 and 100 being the worst and best ranking, respectively (see Supplementary Table 3 for details). To benchmark integration results, we applied ASW using batch labels, with a lower score indicating a lower level of batch effects.

We next sought to assess the efficacy of supervised cell type annotation versus unsupervised clustering using SingleR, relative to their ground truth labels. We took the 8 pancreas samples (Supplementary Table 2) and used each sample as a query and the remaining 7 samples as a reference using their ground truth labels as their annotations.

## Results

### Overview of IBRAP

IBRAP is the most comprehensive scRNA-seq analysis R-based toolkit that is a wrapper for a range of single-cell analytical tools across R and Python, compiled into R. To help users, we produced an installation of 100+ packages and their dependencies, and an S4 class object in R to store large volumes of results. IBRAP enables users to interchange normalisation, integration, and clustering methods throughout pipelines to tailor pipelines to their specific dataset. Benchmarking metrices are also included which can help guide users to select the best pipeline and parameter choices. In comparison to other methods, IBRAP provides the highest number of scRNA-seq-specific normalisation, dimensionality reduction, and clustering methods (Table 1), which enables numerous different pipelines and associated results. Furthermore, several benchmarking metrics were included into our benchmarking function allows for a detailed inspection and comparison of different results. To facilitate the exploration and comparison of the large volume of results IBRAP also provides an ease-of-use Rshiny interface.

#### Rshiny Application

Even with integration benchmarking and a summary of clustering scores, assessing each component through a terminal would be tedious. Thus, we build an ease-of-use Rshiny application called IBRAP Exploration. IBRAP Exploration contains three key areas: interactive reduction visualisation (Supplementary Figure 1A), Benchmarking visualisation (Supplementary Figure 1B), and Gene expression visualisation (Supplementary Figure 1C). Interactive reduction visualisation, contains an interactive plot for displaying reduced dimensions and interchanging cell labels from clustering to important metadata such as, sequencing depth and mitochondrial fractions. Benchmarking visualisation, is used for examining benchmarking metrices that explain clustering and benchmarking performance to aid users in selecting best clustering results. And finally, gene expression visualisation, for dissecting gene expression that may have drove the clustering and can describe cell types contained in clusters. This tool will enable users to interactively and swiftly visualise their results.

### Case Studies

To demonstrate the effectiveness of the tools in IBRAP, we applied all possible pipelines within IBRAP and performed benchmarking on these results to ascertain their effectiveness. We recruited 15 samples (of 21,029 cells) from publicly available sources and 4 simulated datasets (n=4,000) constructed with splatter and accompanied with ground truth cell annotations, covering a range of biological and technical backgrounds (Supplementary Table 2). The sample collection included: (1) 8 pancreas pancreatic samples (cell range: 638-3,605 per sample, 13 cell types; technologies: CELseq, CELseq2, SmartSeq2, Fluidigmc1, inDrop), (2) 7 cell line samples (cell range: 225-3,818 per sample, 3-5 cell lines; technologies: 10x, CELseq2, Dropseq), and (3) 4 simulated samples using splatter [17] (cells: 1,000 per sample, 4-8 cell types). For these studies, we applied all normalisation (SCTransform; Scanpy; Scran), integration (Scanorama; BBKNN; Seurat CCA; Harmony), and clustering methods (louvain; louvainMLR; SLM; Leiden; SC3; Kmeans; PAM).

#### 1. Individual Sample - Unsupervised Analysis

We first applied IBRAP to assess pipeline performance for individual samples. A total of 21 unique pipelines were produced for each sample. After parameter adjustments for the clustering components (e.g., resolution and expected number of clusters), 105 different clustering results were generated per sample, and their performance were assessed and compared (Supplementary Table 4). The average (ARI, NMI, and ASW) values for all pancreas, cell line, and simulation samples were (0.690, 0.747, and 0.559) and (0.804, 0.793, and 0.713), and (0.904, 0.918, and 0.794), respectively (Figure 2A; Supplementary Figure 2). Compared to pancreas samples, cell lines and simulated data often have lower level of cellular complexity, which leads to the requirement of simpler analytical tasks, reflected by the higher average benchmarking scores.

**Figure 2.**
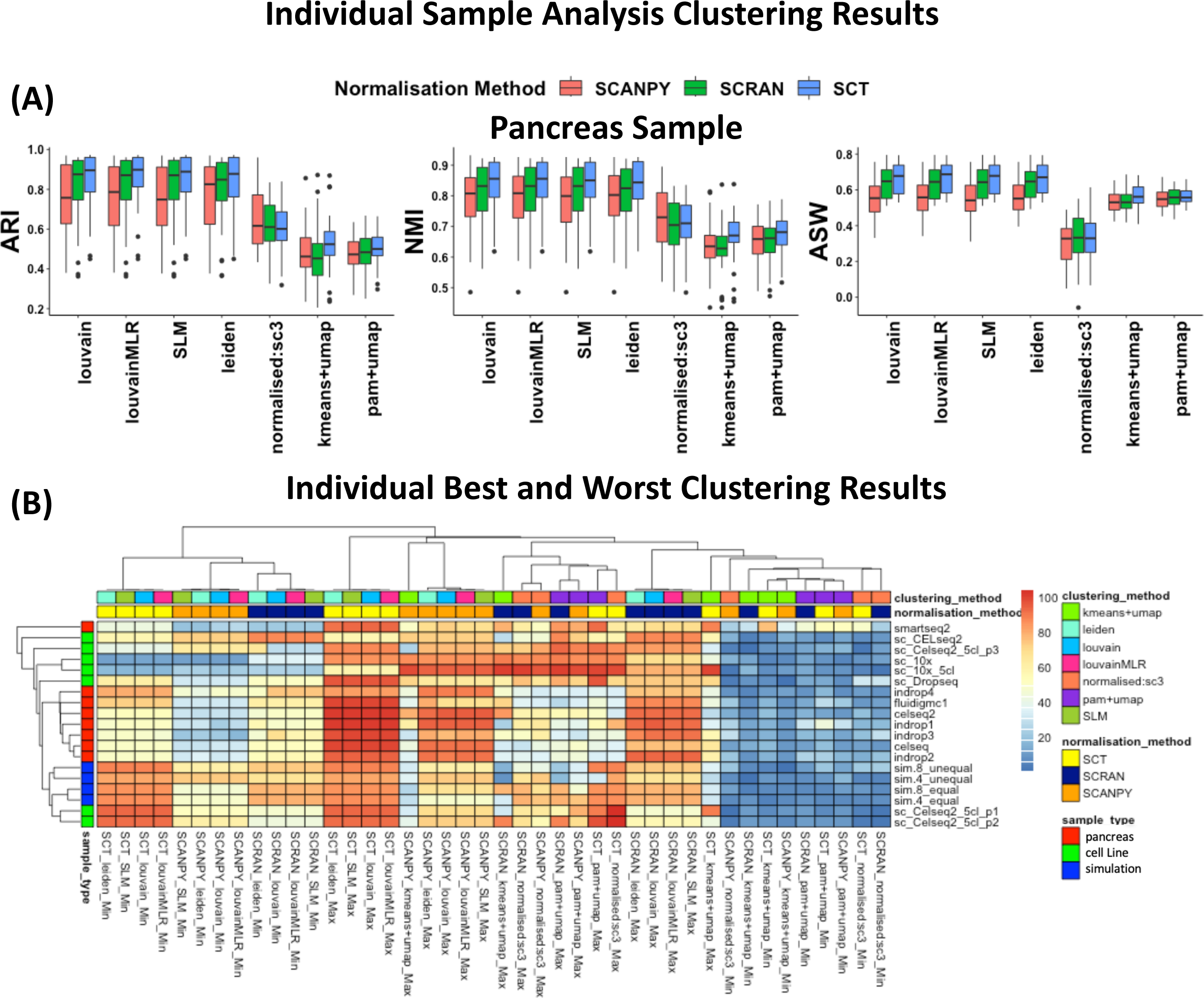
| Individual Sample Analysis Clustering Results. **A** Benchmarking metrices: ARI, NMI, and ASW for pancreas samples. A higher score indicates a better cluster assignment whilst a lower score is less favourable. **B** A heatmap showing the highest and lowest performance scores for each possible pipeline combination (x-axis) compared against each sample (y-axis). A higher (red) score is better whilst a lower (blue) score is worse.

We first assessed how different normalisation techniques affected subsequent clustering performances (Figure 2A&B; Supplementary Figure 2). For pancreas samples, the average ARI, NMI, and ASW values from results generated by SCTransform (ARI=0.720, NMI=0.768, ASW=0.587) were higher than those by Scran (ARI=0.687, NMI=0.742, ASW=0.570) and Scanpy (ARI=0.662, NMI=0.734, ASW=0.520), indicating that pipelines implementing SCTransform produced superior clustering performance on average. Indeed, when identifying the best performing pipelines benchmarked with the ground truth cell labels, pipelines with SCTransform as the normalisation component produced the best clustering performance for 7 out of 8 pancreas pancreatic samples. However, for a single sample (indrop4) Scanpy normalisation yielded the best results combining with graph-based clustering algorithms. For cell line samples, the average ARI, NMI, and ASW values generated by SCTransform (ARI=0.816, NMI=0.805, ASW=0.723) were higher than Scanpy (ARI=0.802, NMI=0.789, ASW=0.702) and Scran (ARI=0.797, NMI=0.784, ASW=0.712). Although SCTransform-derived pipelines still produced the best average ARI and NMI results, only 3 out of 7 cell lines (technologies: 1xDropseq, 2xCELseq2) gained their best clustering performance using SCTransform. Whereas, pipelines consisting of Scran normalisation gave the best performance score for 4 out of 7 samples (technologies: 2 of 10x, 2 of CELseq2) (Supplementary Figure 2; Supplementary Table 4). For simulated samples, ∼48% of the pipelines produced perfect clustering scores regardless of normalisation choices due to simplicity of the samples (Supplementary Figure 2; Supplementary Table 4).

Next, we investigated how clustering algorithm choice affected the overall performance of the pipelines. For pancreas and simulated samples, pipelines consisting of graph-based clustering (Louvain, Louvain with multi-level refinement, SLM and Leiden) produced the best clustering performance in relation to the ground truth cell labels. For cell line samples, there was no consistency in clustering methods incorporated into the highest performing pipelines (Figure 2; Supplementary Figure 2; Supplementary Table 4). It is worth noting that the combination of SCTransform with graph-based clustering methods typically achieved better clustering performances in pancreas samples (Figure 2A&B). Overall, our results of normalisation and clustering methods comparison still demonstrated that there remains no single gold-standard approach for single sample analysis. Therefore, necessitating a tool such as IBRAP, that has the capacity to build the best combinations for the supplied samples.

To further demonstrate the utility of results generated by IBRAP, we compared the best and worst performing pipeline generated for one pancreas pancreatic sample, processed using Smartseq2 (2,394 cells). The best performing pipeline consisted of SCTransform normalisation and SLM (resolution=0.1) clustering (rank score=100, ARI=0.739, NMI=0.803, ASW=0.753) (Figure 3A). Eight clusters were identified compared to 13 cell type ground truth labels (Figure 3B). Major cell types (including: alpha, beta, delta, acinar, ductal, and gamma) were mostly clustered correctly, however minor populations, including: epsilon, mast, macrophage, activated and quiescent stellate, schwann, and endothelial cells were grouped into a single cluster (cluster 7; Figure 3A&B). This is due to very low numbers of these cell types being present in this sample. The worst performing pipeline, consisting of Scanpy normalisation and SLM (resolution=0.4) clustering (score=0, ARI=0.377, NMI=0.581, ASW=0.533) (Figure 3C&D). Sixteen clusters were identified for 13 cell types, major cell types were broken into smaller clusters (i.e., alpha cells into 5 clusters) whilst smaller populations, such as: mast, activated stellate, and endothelial cells were still clustered together. It is worth noting that non-graph-based clustering methods such as, k-means, PAM, and SC3 failed to generate any of the best scores for all pancreas samples (Figure 2).

**Fig 3.**
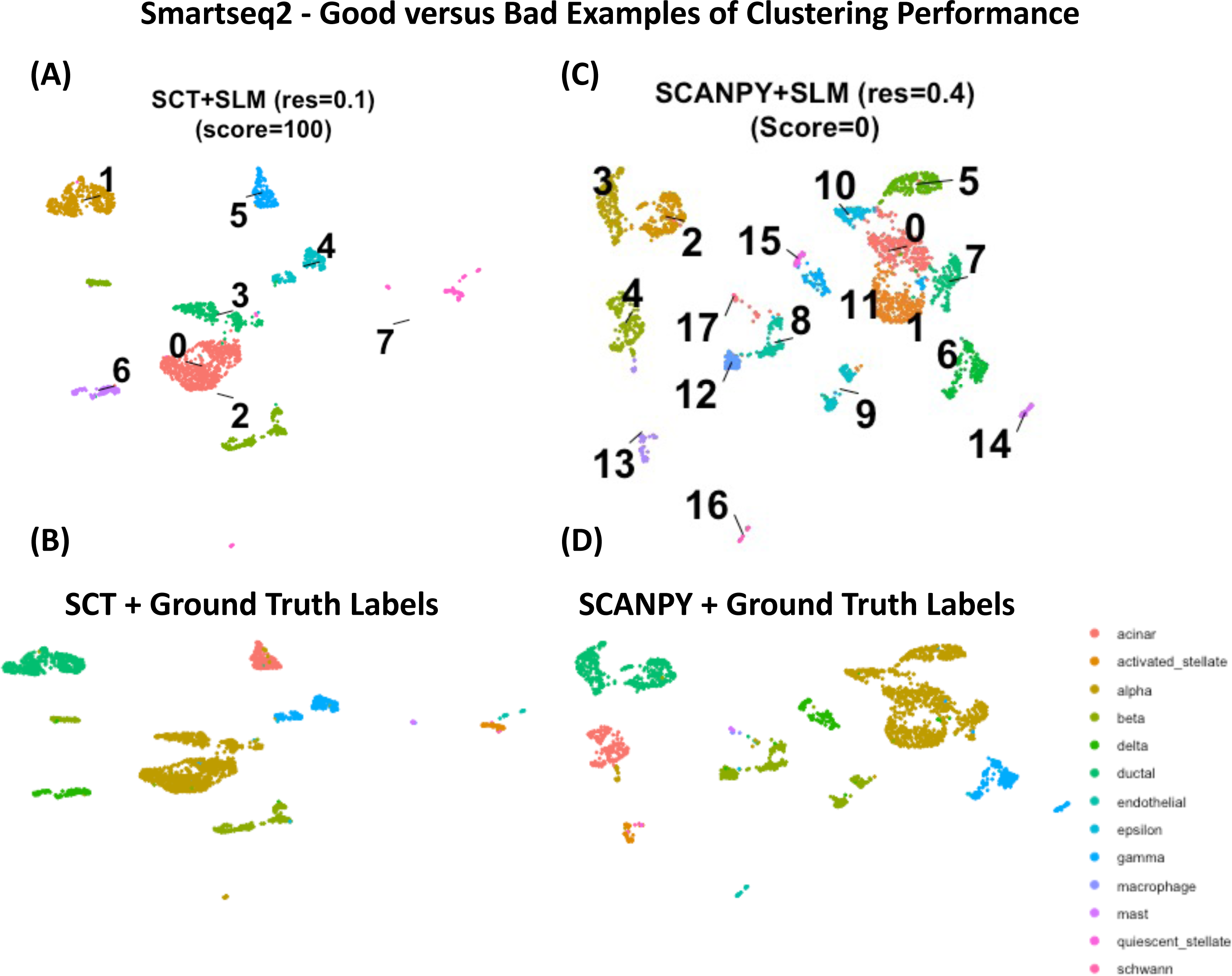
| Individual Sample Cluster Assignments. **A&B** UMAP projection for smartseq2 showing good performing cluster assignments compared against their ground truth. **C&D** UMAP projection for smartseq2 showing bad performing cluster assignments compared against their ground truth.

For the 10x cell line sample (3,918 cells), the best performing pipeline consisted of Scran normalisation and PAM clustering (k=5) (rank score=100, ARI=0.963, NMI=0.942, ASW=0.813) (Supplementary Figure 3A&B). Evidently, all cell line samples benefitted from non-graph-based clustering with a defined number of expected clusters, which reproduced ground truth labels to an almost perfect accuracy. For one of the worst performing pipelines, it consisted of SCTransform normalisation and Louvain (resolution=0.5) clustering (score=15.1, ARI=0.685, NMI=0.664, ASW=0.493) (Supplementary Figure 3C&D). This pipeline resulted in multiple cluster assignment for single cell lines that did not reflect the ground truth labels. Our results further demonstrate that no single pipeline combination is appropriate for all samples, showing the necessity for an integrated tool, such as IBRAP to construct pipelines and compare their performances.

#### 2. Integrating Samples - Unsupervised Analysis

We next evaluated pipelines that required integration methods. We performed 3 multi-sample integration analyses, including, Analysis 1: 8 pancreatic samples (total cells=14,739; technologies=SmartSeq2, CELseq, CELse2, Fluidigmc1, InDrop), Analysis 2: 3 samples with the same cell lines (total cells=1,401; technology=CELseq2, 10x, DropSeq), and Analysis 3: 3 samples that contain 3 of the same cell lines but one sample containing two extra and different cell lines (total cells=4,416; technologies=CELseq2, 10x, Dropseq). A total of 102 unique pipeline combinations were concocted; once we adjusted parameters (resolution or numbers of expected clusters), 1,172 sets of cluster assignments were produced per sample. We then applied our benchmarking metrics to these cell assignments, to compare performances of different pipelines (Supplementary Table 5).

We first assessed the effect that changing normalisation methods would affect the final clustering results (Figure 4). Across all three integration analyses the average (ARI, NMI and ASW) values for SCTransform (blue boxes), Scran (green boxes), and Scanpy (red boxes) was (0.479, 0.548, and 0.443), and (0.462, 0.536, and 0.455), and (0.354, 0.436, and 0.335), respectively. These results show that SCTransform normalisation gained better clustering results across all analyses on average during integration tasks. We then compared the effect normalisation methods had on batch effect removal, we also calculated batch effect removal without using an integration method as a baseline comparison. SCTransform, Scran, and Scanpy normalisation without using batch correction achieved an ASW of 0.196, 0.232, and 0.211, respectively. We then compared the normalisation method with applying an integration method show that SCTransform, Scran, and Scanpy achieved an ASW of 0.038, −0.016, and −0.034, respectively. These results suggest that SCTransform works best at removing most of the batch effect without an integration method whilst Scanpy is the best method to apply before applying an integration. Finally, Scran was the normalisation methods applied in the pipeline (average ASW=-0.17) that was paired with BBKNN that integrated batches the most effectively (Supplementary Figure 4). For Analysis 1, the top performing pipeline (score=100, ARI=0.965, NMI=0.927, ASW=0.636) used Scran normalisation. For Analysis 2, the optimal pipeline (score=100, ARI=0.509, NMI=0.506, ASW=0.794) gained superior cell assignments using Scanpy normalisation. For Analysis 3, The leading pipeline (score=100, ARI=0.860, NMI=0.803, ASW=0.788) produced greater clustering scores from using SCTransform normalisation. These findings suggest that different normalisation methods play an influential role in batch effect removal and must be selected carefully.

**Fig 4.**
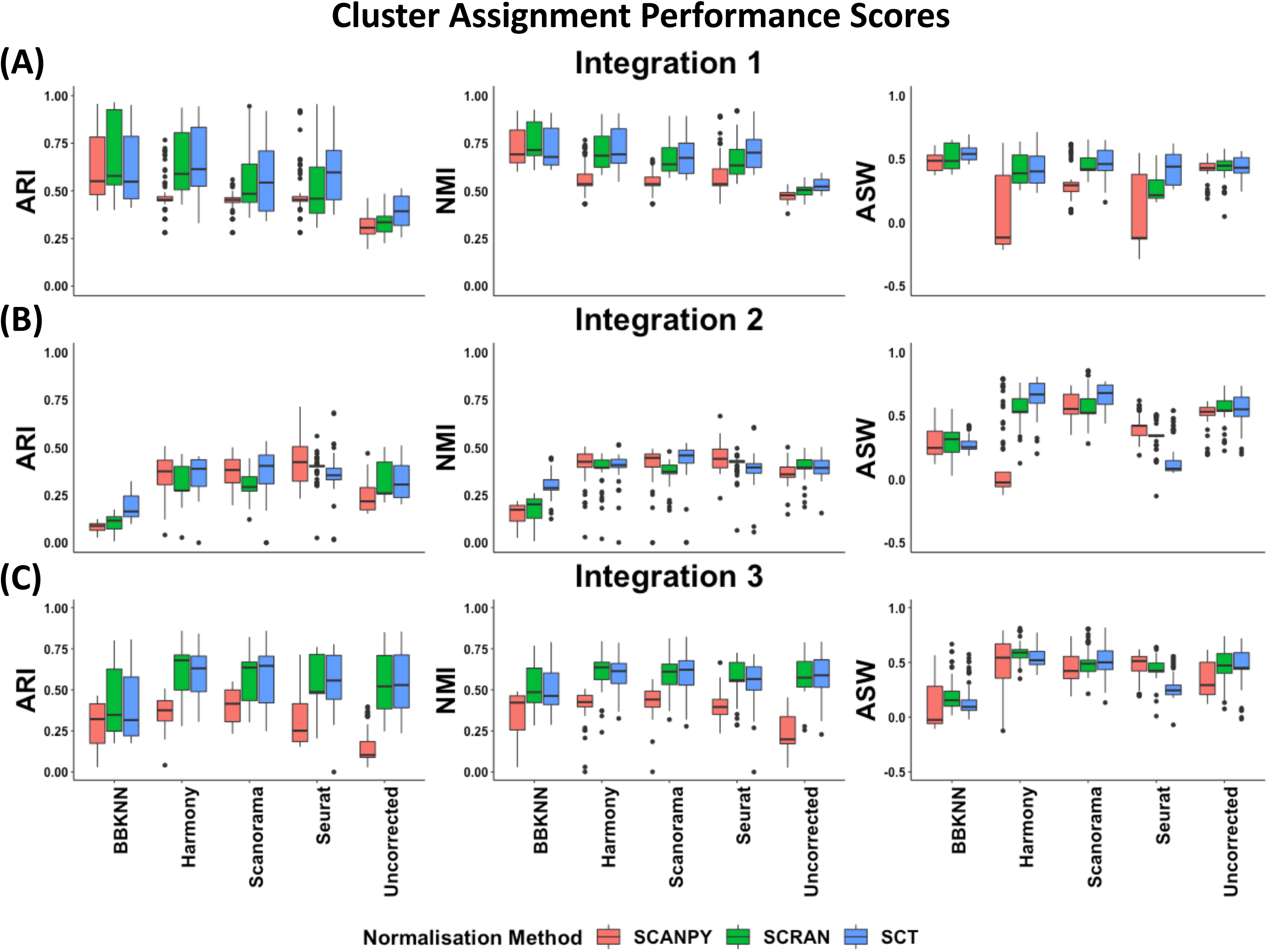
| Benchmarks of multi-sample integration analyses. The analyses included **(A)** Analysis 1: 8 pancreatic samples, **(B)** Analysis 2: 3 samples with the same cell lines, and **(C)** Analysis 3: 3 samples that contain 3 of the same cell lines but one sample containing two extra and different cell lines. Benchmarking matrices, ARI, NMI and ASW, calculated based on ground truth cell labels, were shown for the combinations of normalisation (Scanpy, Scran and SCTransform) and integration methods (BBKNN, Harmony, Scanorama, Seurat CCA and uncorrected), adjusting other clustering methods and parameters.

Next, we investigated the influence of different integration methods (Figure 4A-C separated on the x-axis) on subsequently produced cell annotations in relation to the ground truth labels, and observed differed performances on cluster assignments for uncorrected (an average of ARI=0.365, NMI=0.472, ASW=0.491), Harmony (an average of ARI=0.489, NMI=0.541, ASW=0.383), Scanorama (an average of ARI=0.459, NMI=0.530, ASW=0.515), BBKNN (an average of ARI=0.354, NMI=0.449, ASW= 0.338), and Seurat CCA (an average of ARI=0.486, NMI=0.537, ASW=0.305). These benchmarking matrices showed that Harmony-integrated data resulted in the best cell annotations downstream on average. We next applied ASW to assess the presence of batch effects in the combined datasets and how well integration methods removed them, with a lower ASW value indicating better performance in batch effect. The average ASW for the uncorrected, Harmony, Scanorama, BBKNN, and Seurat CCA was 0.212, 0.100, 0.018, −0.051 and −0.083, respectively (Supplementary Figure 4). These findings suggest that Seurat CCA was the most effective method for batch effect removal on average. However, Seurat CCA did not produce the best overall pipeline for clustering performance for all analyses, suggesting that a higher batch correction does not necessarily lead to the most accurate clustering. In Analysis 1, Scran normalised counts with BBKNN integration generated the best clustering results (score=100, ARI=0.965, NMI=0.927, ASW=0.636) (Supplementary Table 5). Analysis 2 gained its best results using Scanpy normalised counts and Harmony integration (score=100, ARI=0.509, NMI=0.506, ASW=0.794), whilst Analysis 3 obtained the best clustering performance from SCTransform normalisation and Scanorama Integration (score=100, ARI=0.859, NMI=0.803, ASW=0.788),.

Finally, we observed the performance of different clustering performances for integration analyses. On average, non-graph-based clustering (an average of ARI=0.458, NMI=0.506, ASW=0.458) yielded superior clustering results compared to graph-based methods (an average of ARI=0.423, NMI=0.506, ASW=0.379) across the 3 analyses. Indeed, this was also the case when the analyses were broken down individually (i.e., Analysis 1-3). Nonetheless, Analysis 1 gained its best results from Louvain clustering with resolution 0.2 (score=100, ARI=0.965, NMI=0.927, ASW=0.636) whilst Analyses 2 (score=100, ARI=0.509, NMI=0.506, ASW=0.794) and 3 (score=100, ARI=0.859, NMI=0.803, ASW=0.788) gained their best results from PAM clustering with expected clusters as 6, which matches the findings from the individual sample analysis (Figure 4A-C; Supplementary Table 5). These findings are consistent with individual sample analysis, where pancreas samples benefitted from graph-based clustering and cell lines more with non-graph-based methodology.

We then visually inspected a good and bad pipeline performance for Analysis 1. The best clustering results (Supplementary Figure 5A) compared against the ground truth (Supplementary Figure 5B) for pancreatic samples were Scran normalisation, BBKNN integration, and Louvain clustering with the resolution set at 0.2 (score=100, ARI=0.965, NMI=0.927, ASW=0.636). Major cell types were all clustered accurately, although small clusters 8, 7, and 5 contained multiple cell types, including: mast and macrophage, endothelial and schwann, and quiescent and activated stellate, respectively. In contrast, the worst clustering performance (Supplementary Figure 5C) in comparison to ground truth labels (Supplementary Figure 5D) was gained from Scanpy normalisation, Seurat CCA integration, and Leiden clustering with resolution 0.2 (score=0, ARI=0.281, NMI=0.462, ASW=-0.287). Clusters appeared to be spread all across the UMAP and are overlapping each other significantly. This is represented in the ASW value of −0.287 showing that clusters are very poorly separated which has led to poor subsequent clustering. Cell types: ductal, beta, gamma, acinar, endothelial, delta and quiescent stellate have mostly been assigned to cluster 0 with some of the cells been assigned to other clusters. This may be a poor combination of methodology that has led to this, which shows that caution must be taken when integrating new methodology, even when individual methods are among the most widely used, like the case above.

**Fig 5.**
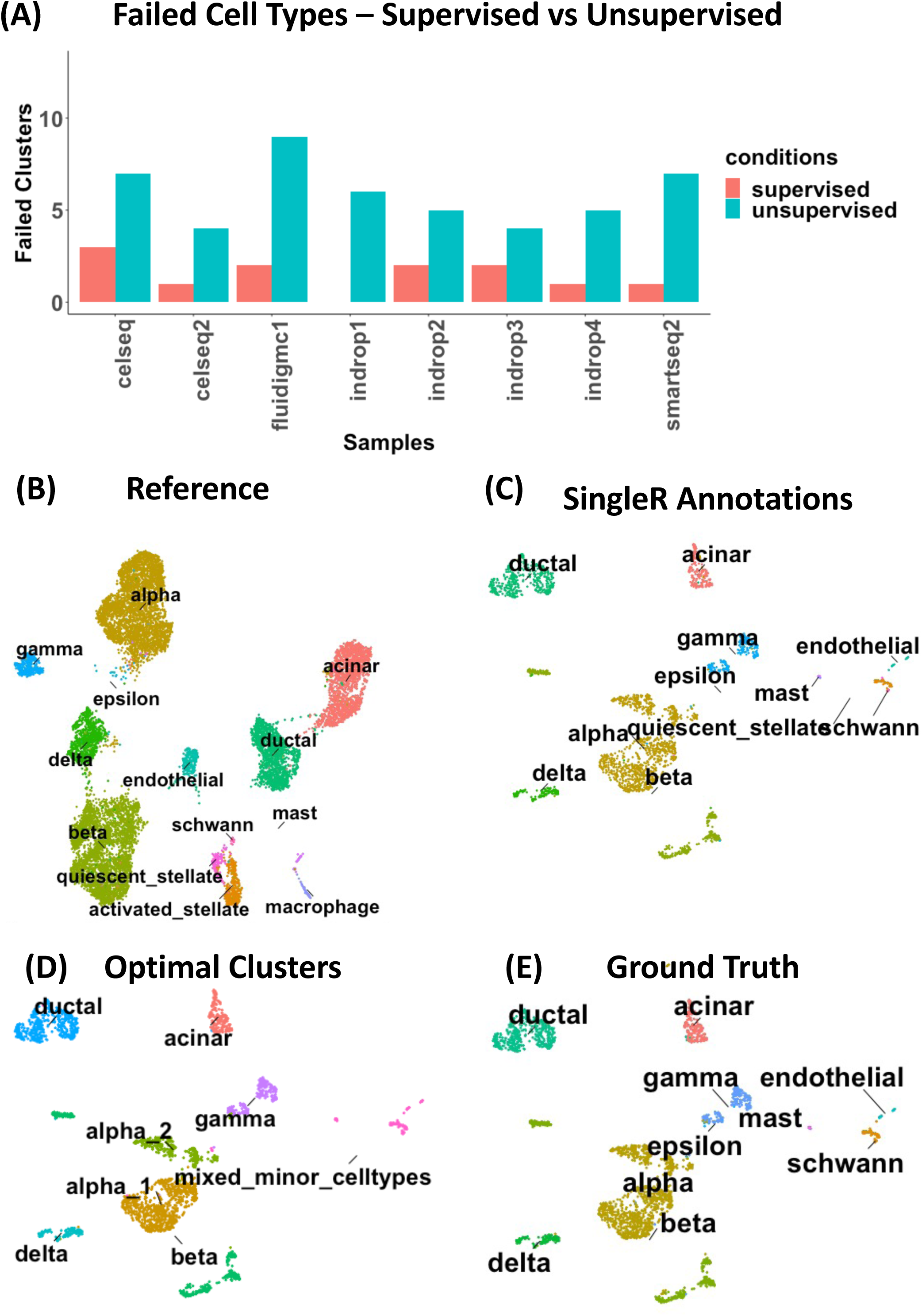
| A comparison between unsupervised and supervised clustering. **A** Bar plot containing the number of cell types that were not captured well during supervised and unsupervised clustering analyses. **B** UMAP projection of the reference datasets (celseq, celseq2, fluidigmc1, indrop1, indrop2, indrop3, indrop4) integrated using BBKNN. **C-E** UMAP projections of the pancreatic samples sequenced using SmartSeq2. **C** Cell labels derived from singleR supervised analysis using a reference dataset in panel B. **D** smartseq2 cells labelled with cluster assignments that generated the highest score. **E** Ground truth labels for comparison to the supervised and unsupervised labelling.

#### 3. Supervised vs Unsupervised Cell Type Annotation

One of the major challenges that scRNA-seq downstream analysis faces is cell annotation, i.e., how to identity each major and minor/rare populations accurately. The standard cell annotation involves unsupervised clustering followed by differential expression and cluster annotation using top differentially expressed genes for each cluster. Cell annotation using reference-based maps (i.e., supervised cell annotation) has, however, been shown to cluster imbalanced cellular proportions (with rare cell types) more effectively compared to unsupervised methods[7]. As both unsupervised and supervised clustering functions were included in IBRAP, we wanted to investigate how the performance varies between the two in cell clustering and annotation, using the 8 pancreatic samples from the individual analysis all with ground truth labels. For supervised clustering, we applied SingleR to each sample individually by using 1 sample from the total of 8 as a query and using the remaining 7 as a reference based on the ground truth labels (Figure 5A). For the unsupervised analysis, we selected the best performing pipeline results generated from the individual sample analysis (Supplementary Table 4). We compared the number of failed clusters between the supervised and unsupervised analyses. We defined a failed cluster with two criteria in relation to the ground truth labels. Firstly, a cell type must reasonably be the dominant cell type in a cluster. Secondly, A cell type cannot be clustered together in similar proportion to other cell types. On average, there were 1.6 and 5.9 failed clusters for the supervised and unsupervised analyses (Figure 5A), respectively. For comparison, we generated several UMAP projections: the seven samples used as a reference for the singleR analysis (Figure 5B), the labels for the supervised analysis for smartseq2 (Figure 5C), the optimal unsupervised clustering labels from the first case study (Figure 5D), and the ground truth cell type labels for smartseq2 (Figure 5E). The supervised analysis (Figure 5C) identified all cell types except for activated stellate cells. However, during the optimal unsupervised analysis, rarer populations (quiescent and activated stellate, mast, macrophage, etc.) were merged into the same cluster (labelled mixed_minor_celltypes in Figure 5D). These results suggest that rarer populations are more readily captured in supervised analyses compared to unsupervised methods in this case study. This is because the reference map based on 7 samples (consisting of 12,496 cells in total) had decent enough resolution to differentiate rarer populations, such as quiescent *vs.* activated stellate cells (Figure 5B). These differential expression profiles of cells provided a prior knowledge to separate cells even if the differences are subtle in a single query sample of 2,395 cells in this example (smartseq2, Figure 5C). However, these subtle differences among related but different cell types could not be resolved by unsupervised clustering analysis alone, especially when the cell numbers were low. It is worth noting that supervised methods are heavily constrained by the quality and the bandwidth of cell types in the supplied reference. If the reference is low in resolution, it would be very difficult to identify rare and novel populations as previously described [7].

## Discussion

Single-cell studies are becoming more popular in the understanding of cellular heterogeneity and interaction in health and disease. Subsequently, a large body of dry-lab analytical methods are becoming available for bioinformaticians to use. Our results demonstrate that different pipelines yielded varying efficacy according to the combination of tools used throughout the pipeline and the input data, which supports previous findings [2,4–12]. Therefore, we demonstrated the requirement for a tool that facilitates different combinations of methods to enhance sensitivity and specificity of data outcomes.

IBRAP is a wrapper package that is fit for this purpose and an attractive alternative for bioinformaticians to use to expand their pipeline possibilities beyond the constraints of current analytical packages for scRNA-seq. To aid users, we have included benchmarking metrices that can help to guide users to ascertain the effectiveness of pipelines. Thus, IBRAP can enhance scRNA-seq pipelines and encourage bioinformaticians to explore other pipeline combinations and generate more accurate results, which may not be possible using single package tools/pipelines.

There are a number of limitations regarding IBRAP which are common to all scRNA-seq analytical packages such as, not including FASTQ alignment to a reference genome. Another downfall is the ability to infer RNA velocity from spliced/unspliced matrix counts and differential abundance tools. However, IBRAP does provides a bespoke tool to enable users to tailor their downstream analyses specifically for their input data and to benchmark their results. IBRAP – like other scRNA-seq platforms – allows for cell filtration, normalisation, quality control, dimensionality reduction, integration, clustering, differential expression analysis, and trajectory analysis. IBRAP is further limited by the requirement to maintain the package alongside dependency updates; this is somewhat overcome by IBRAP being available as a docker image.

In our case studies as we benchmarked pipelines that were generated for individual sample and integration-based analyses based on 19 samples from different biological conditions and technical platforms, we can provide some guidance for future studies. During the normalisation stage, we found that on average SCTransform and Scran led to the best clustering results when analysing samples individually and integrating them, respectively. In terms of integration, we discovered that Harmony led to the most accurate cell type label annotations on average, but Seurat CCA eliminated the most batch effects on average. However, we believe the reason that the most batch effect removal did not correlate with the best cell type labels is likely due to Seurat CCA over-correcting data[10]. Finally, we identified a trend in clustering algorithms, where pancreas samples benefitted from graph-based methods and cell lines gained their best results from non-graph-based methods across all samples. For example, SCTransform normalisation and Louvain clustering performed well for most pancreas samples, whilst SCTransform and PAM clustering performed well for most cell line samples.

We also demonstrated in our case study that unsupervised clustering methods innately clustered rarer and transcriptional related cell populations together or into larger cell populations, especially when their cell numbers were very low. Whereas supervised (reference-based) methods were more capable of identifying these minor populations together with other cellular compositions even with low cell numbers. Our results further highlighted the advantages of reference-based cell annotation; however, such analysis heavily relies on the resolution and bandwidth of cell types annotated within the reference. Since there has been a huge body of scRNA-seq data sets available from a wide range of health and disease settings, exemplified by Single Cell Portal (https://singlecell.broadinstitute.org/single_cell) and the human cell atlas [32], such rich resources have made it possible to construct accurate and high-resolution references across many biomedical settings, and this has become the priority of current scRNA-seq research, as shown by recent efforts with celltypist [33], HuBMAP [18], and CAP, a Human Cell Atlas led reference map initiative. However, high quality reference maps are still lacking in many settings, and more work is needed with valid underlying research questions.

It is worth noting that when assembling a range of data sets to build a reference or comparable counterparts, unsupervised analysis and manual cell annotation has to be conducted first, thus, it is important to use homogenised pipelines to analyse all the data sets together. IBRAP’s components, specifically pipeline benchmarking and integration methods, were carefully selected to be scalable to reference and Atlas-sized projects, followed by supervised clustering to analyse new samples also formed as part of IBRAP. Therefore, IBRAP presents the community with an attractive, valid and easy-to-use analytic platform, not only for single sample analysis, but also for multi-sample and cross study integration.

## Author contributions

C. K. built the tool, co-designed, wrote the tutorials, gathered the data, and analysed it. J. W. conceived and supervised the project. C. K. and J. W. wrote the manuscript. F. K. co-designed, tested the tools and co-wrote the tutorial. All authors critically reviewed the manuscript.

## Code Availability

IBRAPs code is publicly available and can be found on GitHub alongside any appropriate tutorials - https://github.com/connorhknight/IBRAP.

## Supplementary Figures and Tables

**Supplementary Fig 1.**
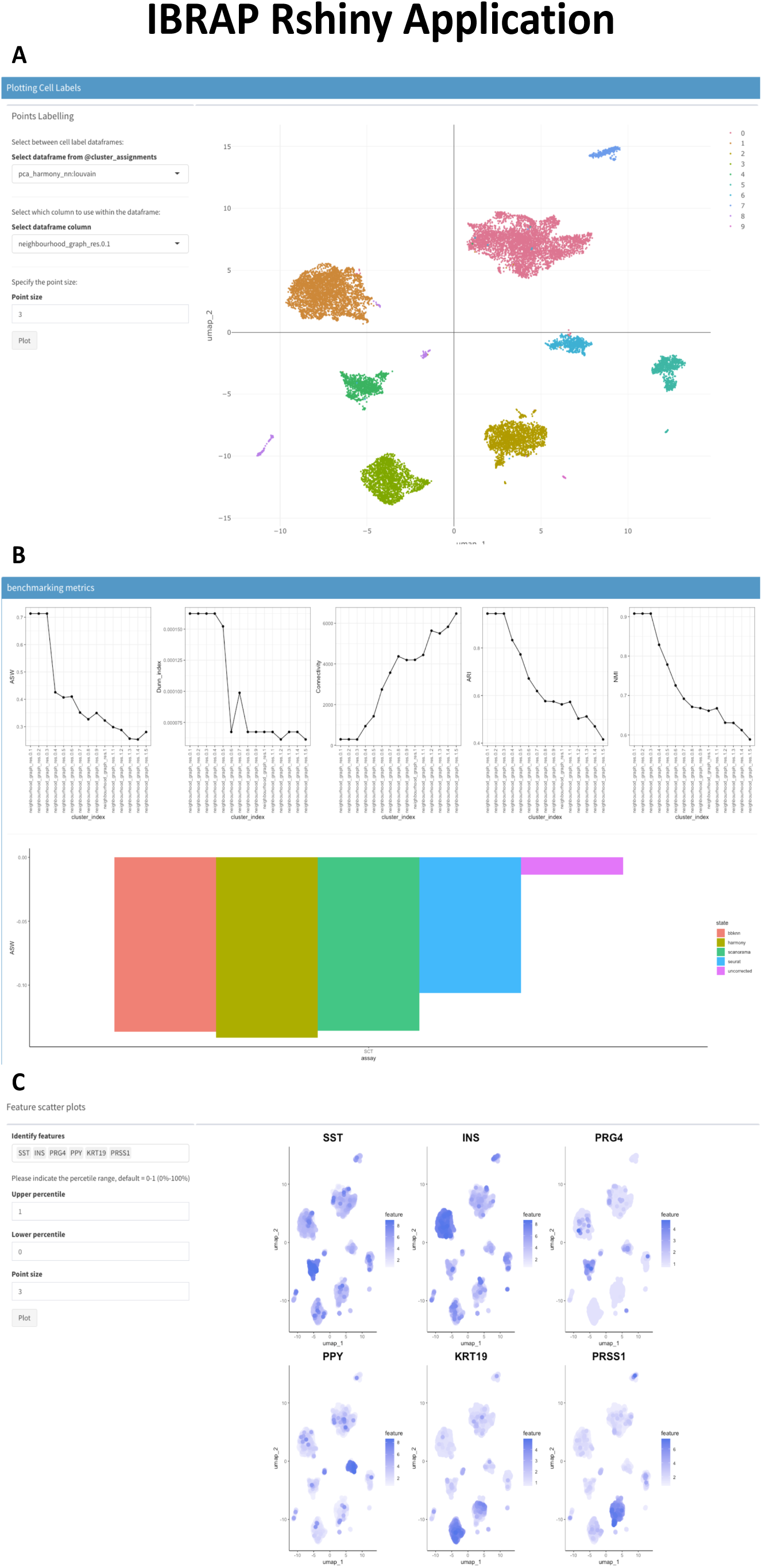
**A** Dimensionality reduction can be interactively (plotly) plotted and labelled with the user’s choice; point sizes can be adjusted accordingly and certain clusters can be omitted. **B** Benchmarking metrices – when available – are displayed below the interactive plots to guide users in their plotting choices. Clustering metrices are displayed the line plots whilst batch effect metrices are shown below this. **C** To dissect the biological information that drove the cluster, feature plots (and other gene-based plots) can be displayed.

**Supplementary Fig 2.**
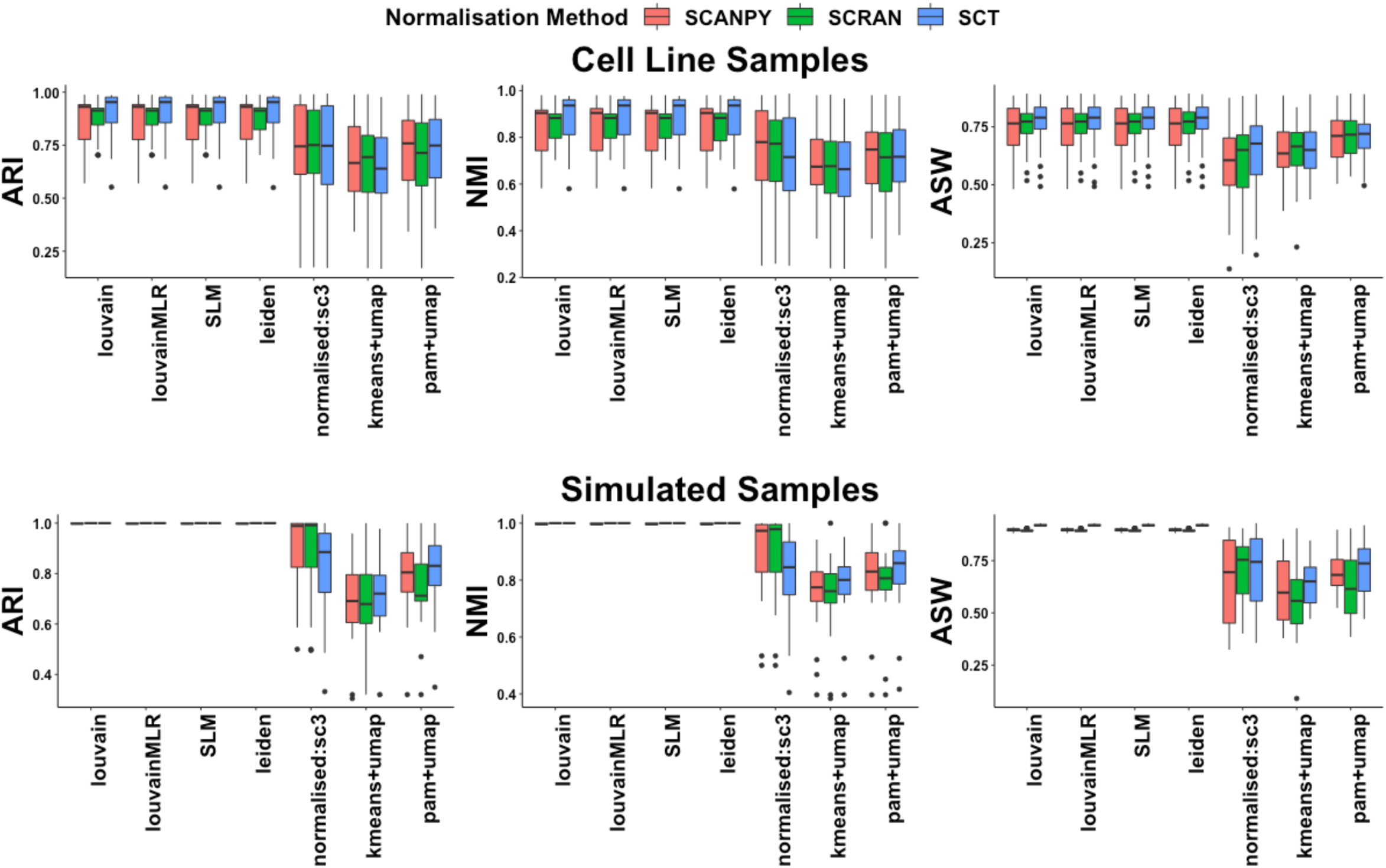
Benchmarking metrices: ASW, NMI, and ASW for cell line and simulated samples. A higher score indicates a better cluster assignment whilst a lower score is less favourable.

**Supplementary Fig 3.**
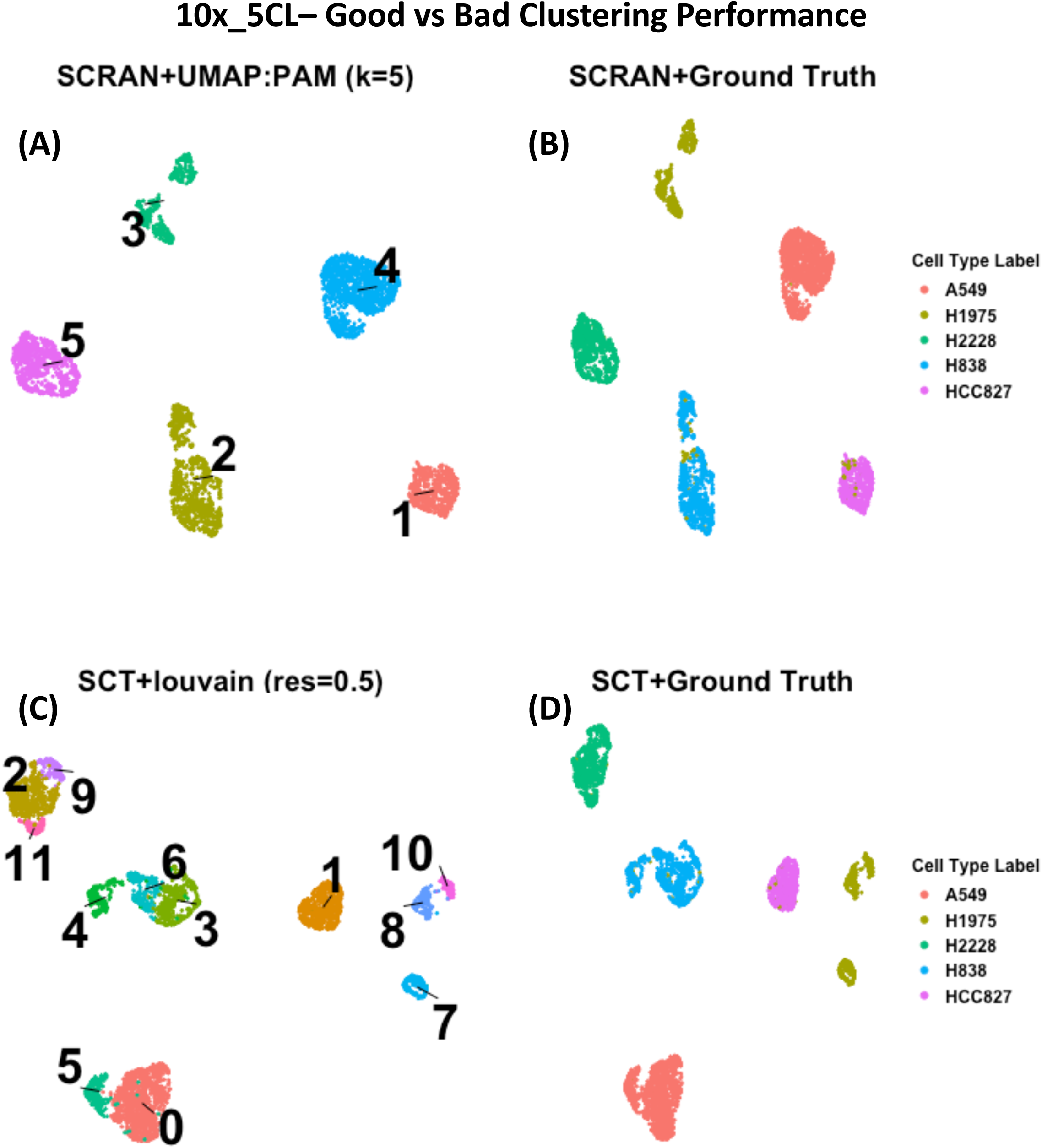
UMAP projections containing good and bad cases for 10x 5 cell line sample. (A) a projection of good clustering using the pipeline specified in the title. (B) The ground truth in relation to panel A. (C) a projection of bad clustering using the pipeline specified in the title. (D) The ground truth in relation to panel C.

**Supplementary Fig 4.**
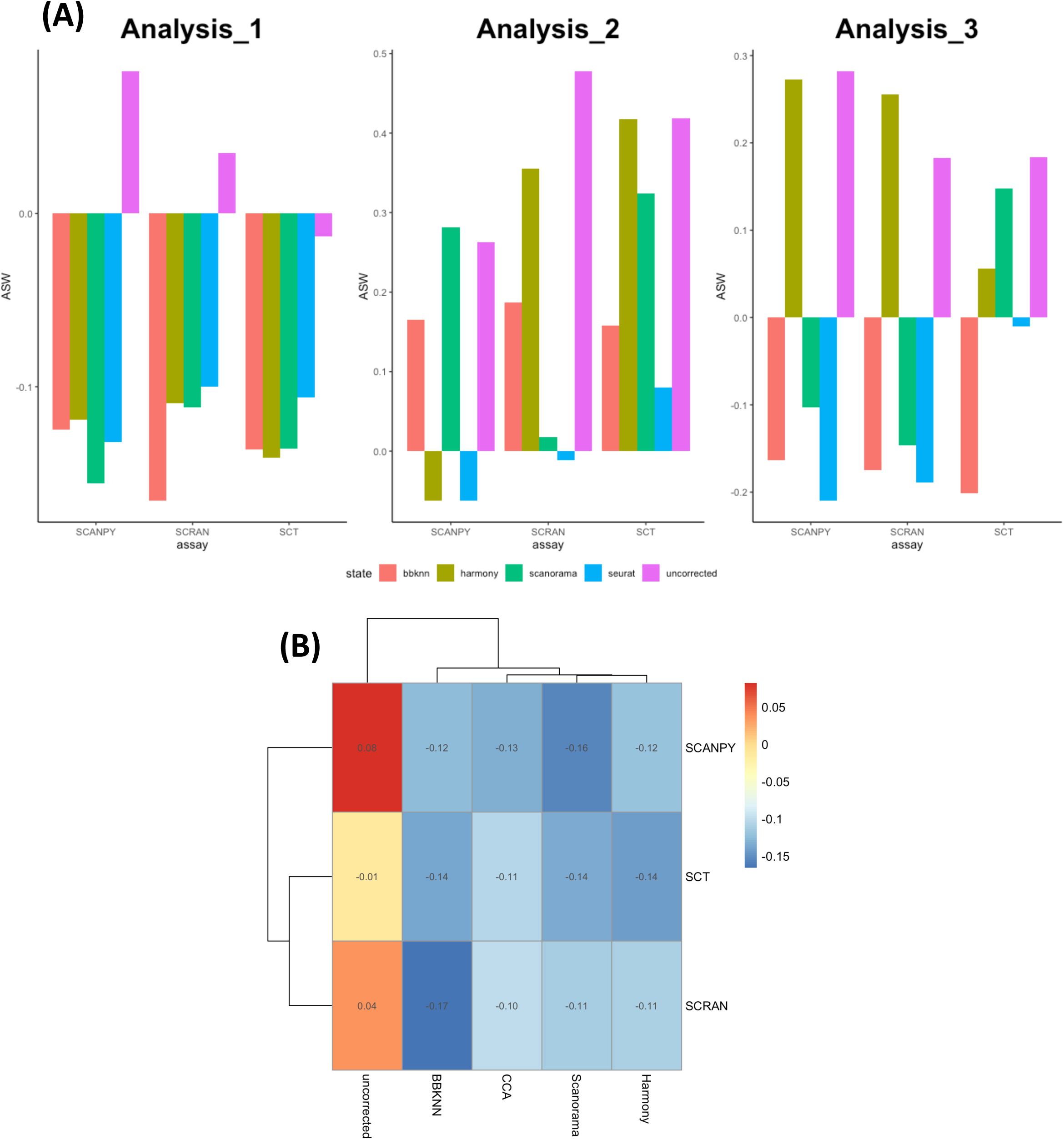
(A) Bar plots showing the batch ASW score between batches before and after correction. A higher value indicates that a higher degree of batch effect is present between batches, whereas a lower value means there is less distance between batches and thus, less batch effects. (B) the average of the normalisation and integration method performance across all analyses

**Supplementary Fig 5.**
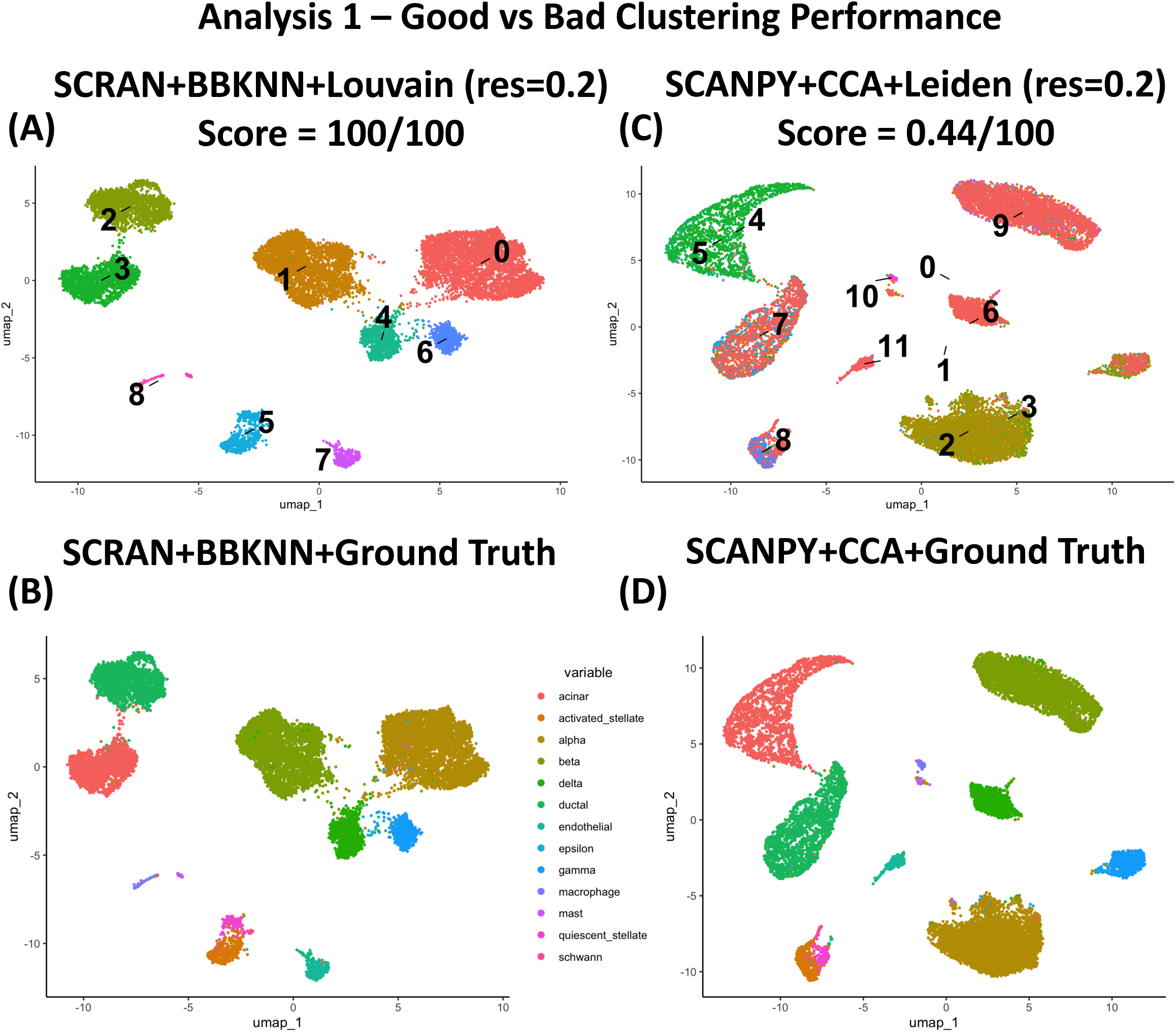
UMAP projections containing good and bad cases for Analysis 1 during multi-sample analyses. (A) a projection of good clustering using the pipeline specified in the title. (B) The ground truth in relation to panel A. (C) a projection of bad clustering using the pipeline specified in the title. (D) The ground truth in relation to panel C.

**Supplementary Fig 6.**
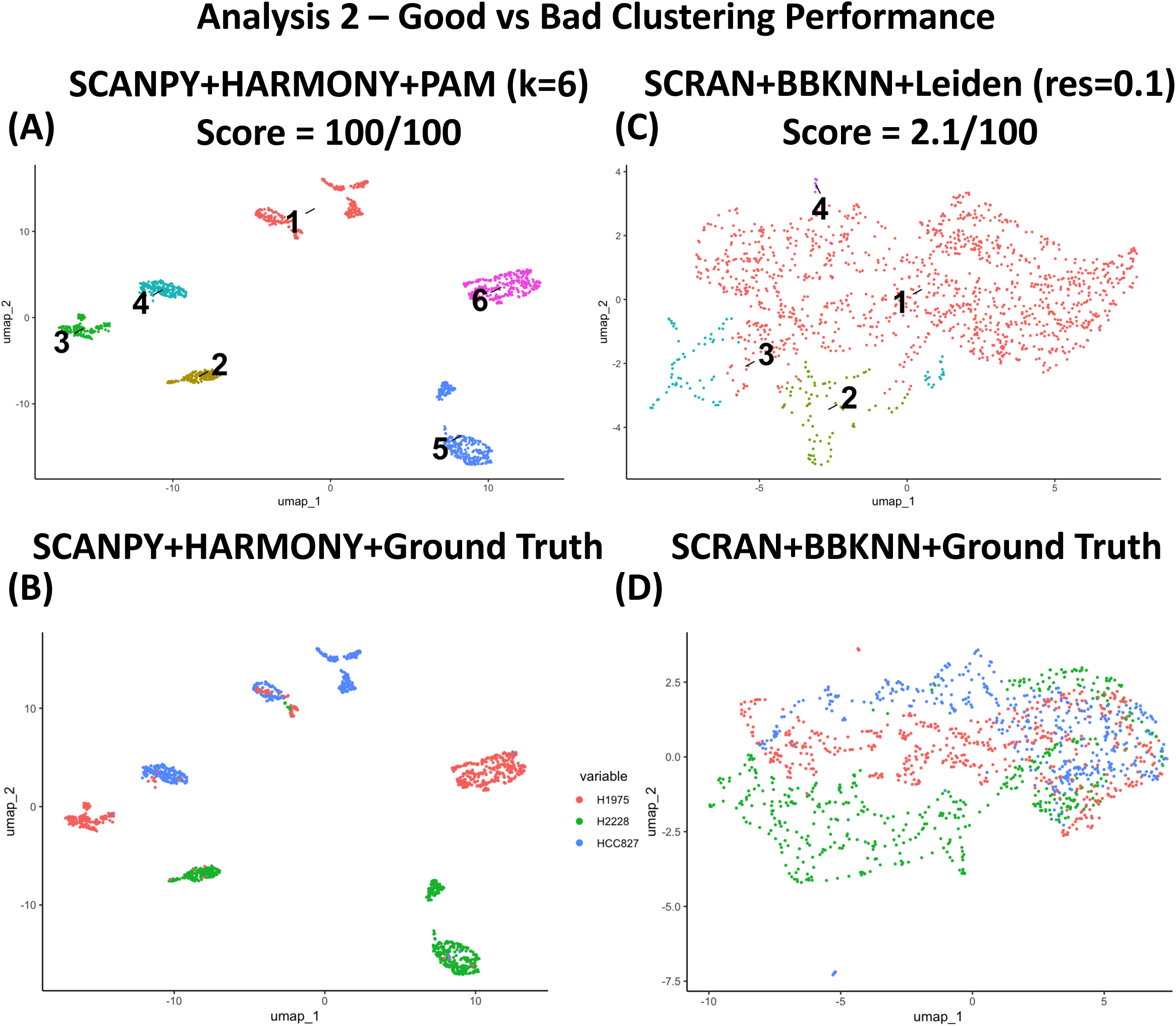
UMAP projections containing good and bad cases for Analysis 2 during multi-sample analyses. (A) a projection of good clustering using the pipeline specified in the title. (B) The ground truth in relation to panel A. (C) a projection of bad clustering using the pipeline specified in the title. (D) The ground truth in relation to panel C.

**Supplementary Fig 7.**
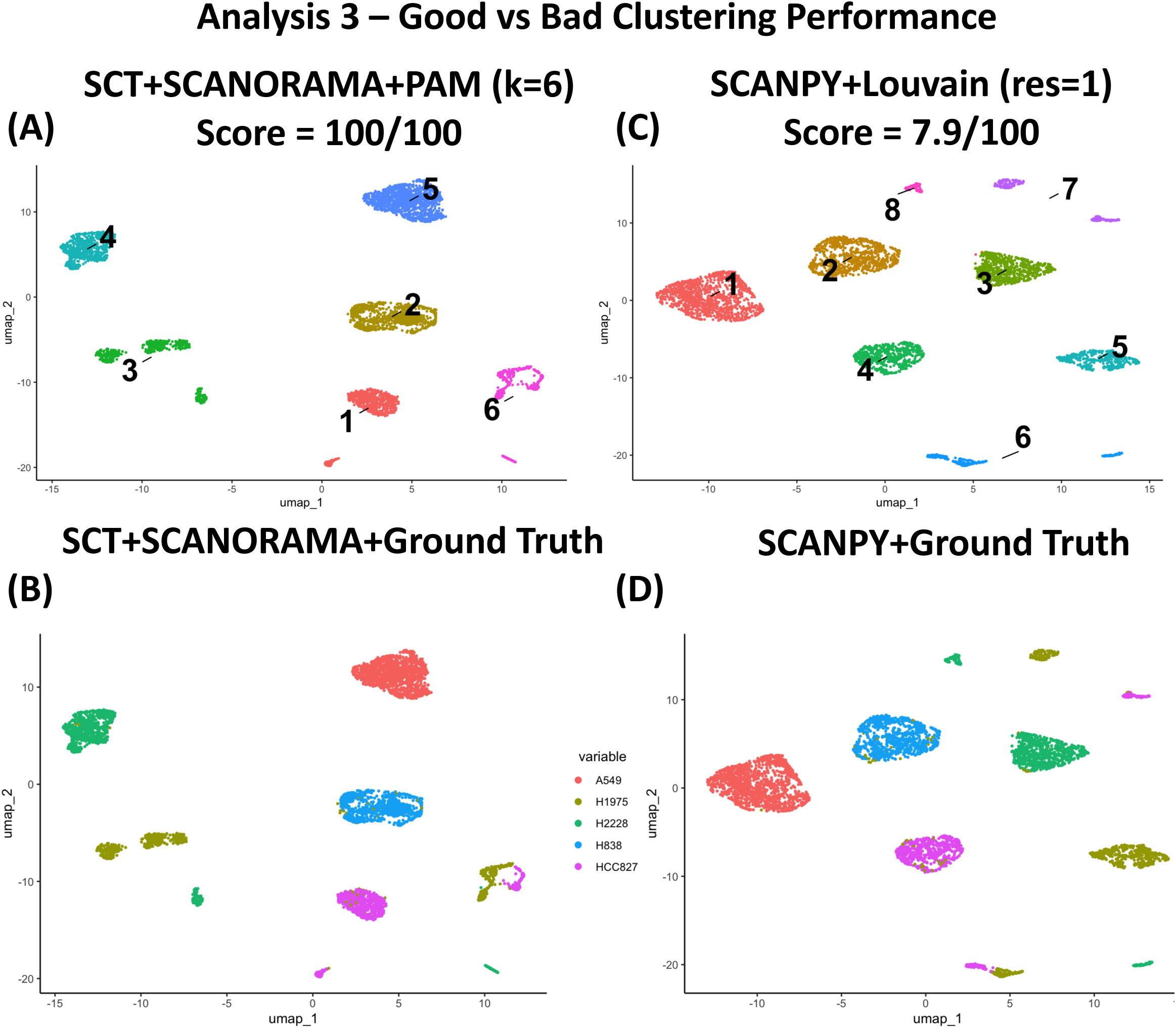
UMAP projections containing good and bad cases for Analysis 3 during multi-sample analyses. (A) a projection of good clustering using the pipeline specified in the title. (B) The ground truth in relation to panel A. (C) a projection of bad clustering using the pipeline specified in the title. (D) The ground truth in relation to panel C.

**Supplementary Fig 8.**
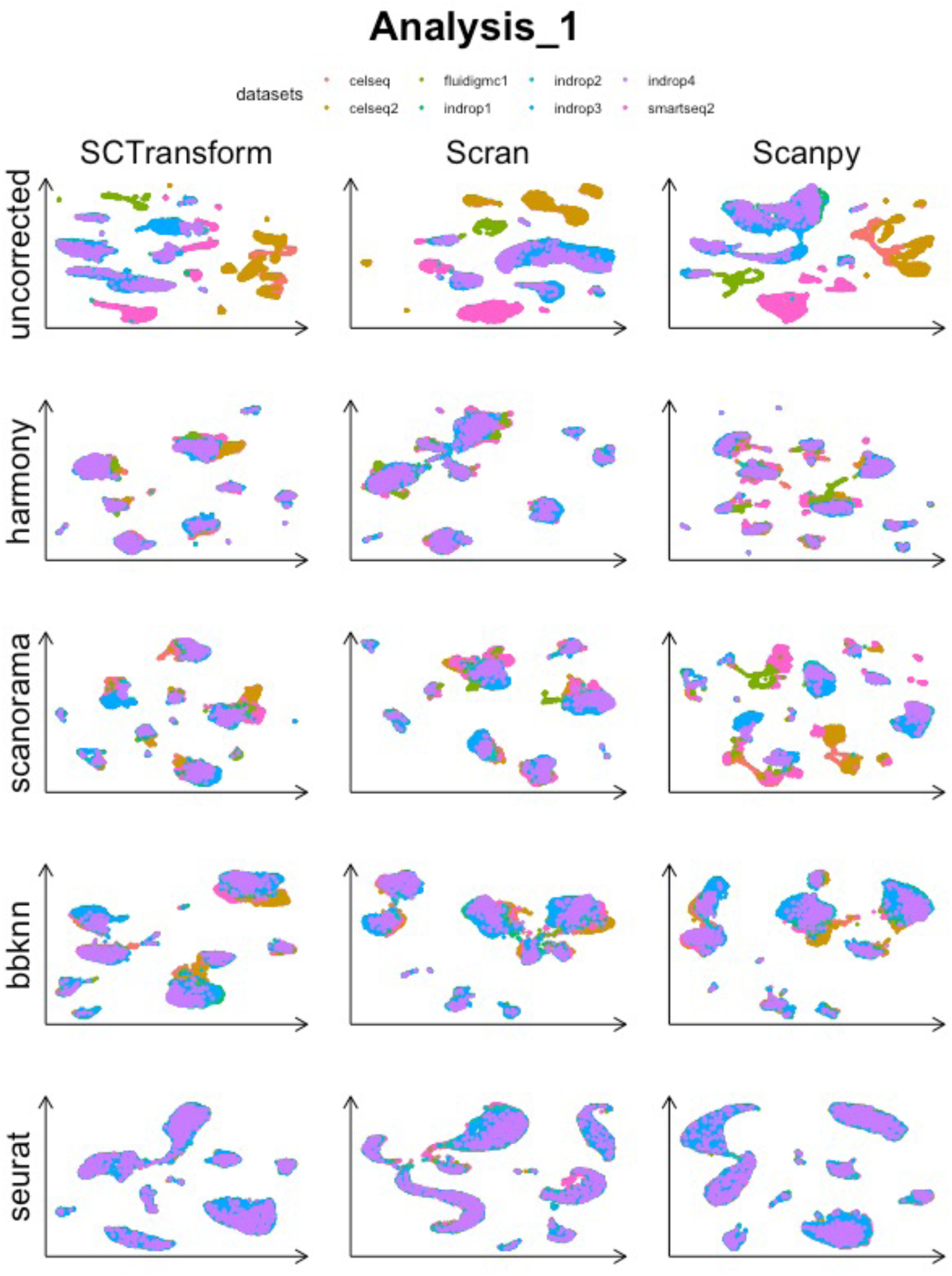
UMAP projections labelled with the batches before and after integration methods for Analysis 1.

**Supplementary Fig 9.**
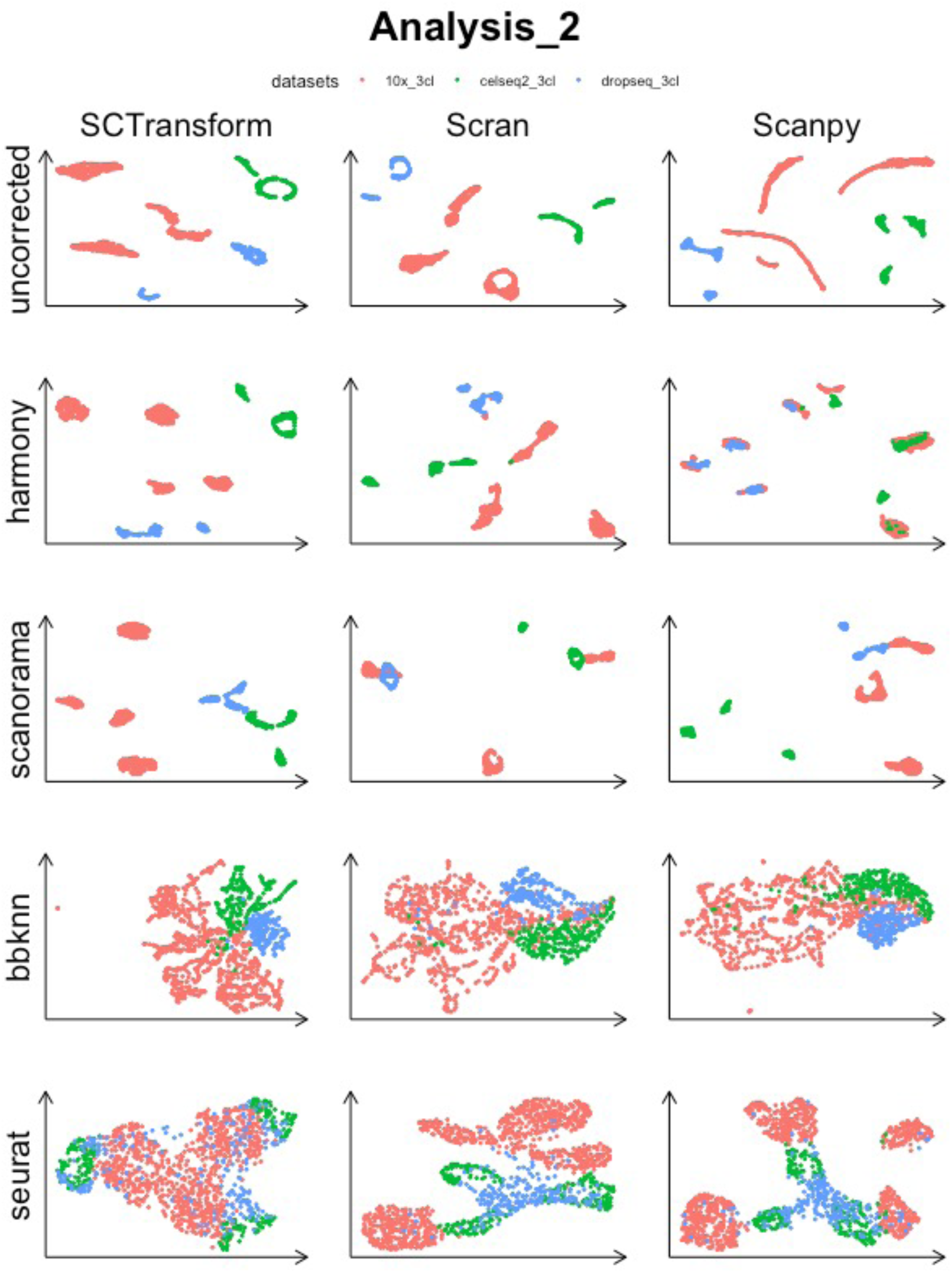
UMAP projections labelled with the batches before and after integration methods for Analysis 2.

**Supplementary Fig 10.**
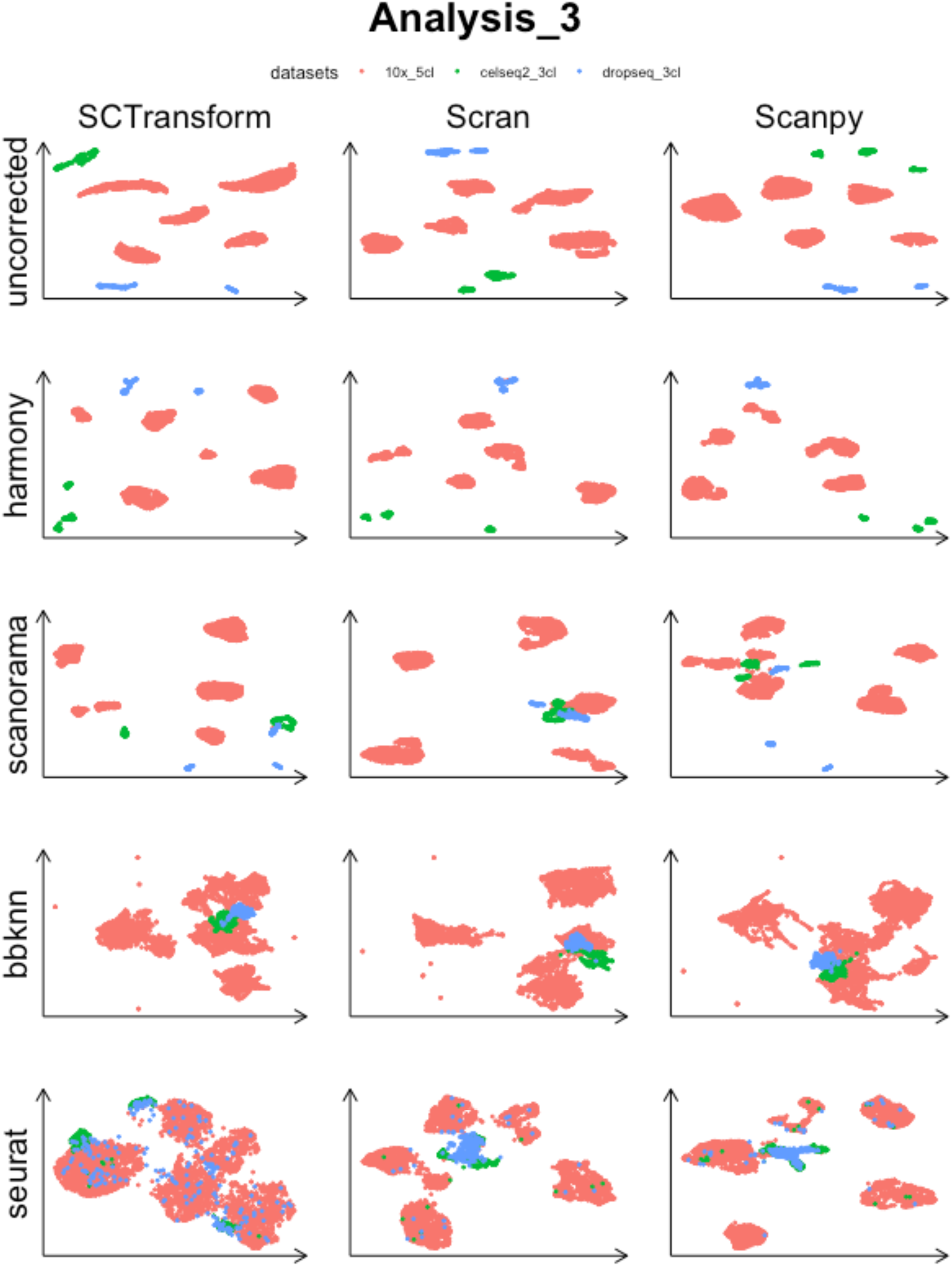
UMAP projections labelled with the batches before and after integration methods for Analysis 3.

**Supplementary Table 1 | Tools Integrated into IBRAP.** A table of tools contained inside of IBRAP separated by the stage where they are applied.

**Supplementary Table 2 | Publicly Available Datasets.** 19 publicly available datasets that have been applied throughout cases studies 1-3. There are 8 human pancreatic tissue samples and 7 mixed cell line samples that were sequenced using a range of library preparation methods. and 4 simulated samples generated using Splatter R package.

**Supplementary Table 3 | Benchmarking Metrices Integrated into IBRAP.**

**Supplementary Table 4 | Case Study #1 Pipeline Components and Benchmarking Scores.** 19 individual samples including pancreas, mixed cell line, and simulations with and 105 clustering results per sample. Here we have applied SCTransform, Scran, and Scanpy for normalisation methods and then then Louvain, LouvainMLR, SLM, Leiden, SC3, PAM, and *k*-means (with a range of expected clusters) to either the normalised counts or reduced dimensions (PCA). Each row in this data frame corresponds to individual normalisation + clustering + adjusted expected clusters per sample.

**Supplementary Table 5 | Case Study #2 Pipeline Components and Benchmarking Scores.** 3 integration tasks were performed using (pancreas) 8 pancreas samples (equal) 3 samples with the same mixture of cell lines (unequal) 3 samples containing some mutual and some unique cell lines, yielding 1,172 clustering results per analysis. We have applied SCTransform, Scran, and Scanpy for normalisation methods and then Harmony, BBKNN, Seurat CCA, and Scanorama and then Louvain, LouvainMLR, SLM, Leiden, SC3, PAM, and *k*-means (with a range of expected clusters) to the integrated reductions. Each row in this data frame corresponds to individual normalisation + integration + clustering + adjusted expected clusters per sample.

